# Circadian rhythms of early afterdepolarizations and ventricular arrhythmias in a cardiomyocyte model

**DOI:** 10.1101/2020.09.13.295436

**Authors:** Casey O. Diekman, Ning Wei

## Abstract

Sudden cardiac arrest is a malfunction of the heart’s electrical system, typically caused by ventricular arrhythmias, that can lead to sudden cardiac death (SCD) within minutes. Epidemiological studies have shown that SCD and ventricular arrhythmias are more likely to occur in the morning than in the evening, and laboratory studies indicate that these daily rhythms in adverse cardiovascular events are at least partially under the control of the endogenous circadian timekeeping system. However, the biophysical mechanisms linking molecular circadian clocks to cardiac arrhythmogenesis are not fully understood. Recent experiments have shown that L-type calcium channels exhibit circadian rhythms in both expression and function in guinea pig ventricular cardiomyocytes. We developed an electrophysiological model of these cells to simulate the effect of circadian variation in L-type calcium conductance. We found that there is a circadian pattern in the occurrence of early afterdepolarizations (EADs), which are abnormal depolarizations during the repolarization phase of a cardiac action potential that can trigger fatal ventricular arrhythmias. Specifically, the model produces EADs in the morning but not at other times of day. We show that the model exhibits a codimension-2 Takens-Bogdanov bifurcation that serves as an organizing center for different types of EAD dynamics. We also simulated a 2-D spatial version of this model across a circadian cycle. We found that there is a circadian pattern in the breakup of spiral waves, which represents ventricular fibrillation in cardiac tissue. Specifically, the model produces spiral wave breakup in the morning but not in the evening. Our study is the first to propose a link between circadian rhythms and EAD formation and suggests that the efficacy of drugs targeting EAD-mediated arrhythmias may depend on the time of day that they are administered.

**Significance Statement:** Why are life-threatening cardiac arrhythmias more likely to occur in the morning than in the evening? The electrical properties of the heart exhibit daily rhythms due to molecular circadian clocks within cardiomyocytes. Our computational model of ventricular myocytes shows that clock-controlled expression of a voltage-gated calcium ion channel leads to early afterdepolarizations (EADs) at certain times of the day. EADs, in which the membrane potential of a cardiomyocyte depolarizes a second time before fully repolarizing, can trigger arrhythmias. To our knowledge, this is the first study linking the circadian clock to EAD formation. Our results suggest that the efficacy of anti-arrhythmic medications targeting this calcium ion channel may depend on the time of day the drug is taken.

## Introduction

Sudden cardiac arrest (SCA) is the most common single cause of natural death in the United States [62]. Distinct from a heart attack, SCA occurs when the electrical system of the heart malfunctions, often without prior symptoms. It is usually caused by arrhythmias such as ventricular tachycardia and ventricular fibrillation. These abnormally fast and irregular heartbeats do not pump blood properly and can cause sudden cardiac death (SCD) within minutes if emergency treatment is not begun immediately [61].

The risk of sudden cardiac arrest is not constant throughout the day. SCD is more likely to occur in the morning than in the evening [47, 72]. Ventricular tachyarrhythmias also exhibit a diurnal rhythm with a peak in the morning [32, 56]. The biophysical mechanisms underlying these daily rhythms in adverse cardiovascular events are not fully understood. The master circadian (∼24-hour) pacemaker in the hypothalamus, the suprachiasmatic nucleus (SCN), influences a variety a cardiovascular phenomenon by coordinating daily rhythms in the release of hormones and other circulating molecules. Recently, it has been demonstrated that circadian clocks within heart muscle cells (cardiomyocytes) also regulate rhythms in cardiac electrophysiology [3].

These intracellular circadian clocks are comprised of transcriptional and translational feedback loops that lead to ∼24-hour rhythms in gene expression. In mice, cardiac ion channel expression and myocardial repolarization are under the control of a clock-dependent oscillator that regulates potassium channel-interacting protein 2 (KChIP2), a subunit required for generating the transient outward potassium current I_to_ [29]. Reduced I_to_ amplitude has arrhythmogenic consequences, perhaps due to lengthened QT (repolarization and depolarization) intervals, and may contribute to sudden death in the early stages of human heart failure [25]. The effect of circadian variation in potassium current on action potential (AP) duration and QT interval has been studied using mathematical models of murine, guinea pig, and human myocytes [21, 60]. Recent experiments in guinea pig myocytes have shown that L-type calcium current (I_CaL_) is under circadian control as well, possibly through the PI3K-Akt signaling pathway [9]. How QT interval is affected by circadian oscillations in the concentration of sodium, potassium, and calcium ions in plasma was also studied in biophysically detailed models of human left ventricular cardiomyocytes [20].

In addition to lengthened QT intervals, the presence of early afterdepolarizations (EADs) is also associated with the development of ventricular arrhythmias [71]. EADs are voltage deflections that occur before full repolarization of the membrane potential during an AP. Extensive modeling of EADs has been performed to understand the ionic and dynamical mechanisms involved in the generation of EADs in isolated cells and their spatial propagation in cardiac tissue [33, 73, 77, 68, 34, 70]. At a basic level, EADs result from reduced repolarization reserve due to reduced outward potassium currents or elevated inward calcium currents [53]. Thus, circadian variation in these currents could render myocytes more vulnerable to EADs at certain times of day, and play a role in the observed circadian profile of ventricular arrhythmias and SCDs.

In this paper, we use biophysical modeling and dynamical systems analysis to study how circadian variation in ionic conductances affects EAD generation. First, we fit a conductance-based model to published electrophysiological data from guinea pig ventricular myocytes at two circadian time points. We then perform simulations of single-cell and 2-D spatial domain versions of the model across a circadian cycle. In the single-cell model, we find that EADs occur in the morning but not at other times of day. In the spatial model, we observe that spiral wave breakup, a phenomenon associated with ventricular arrhythmias in cardiac tissue, occurs in the morning but not in the evening. We also show that the single-cell model exhibits a codimension-2 Takens-Bogdanov bifurcation, which can serve as an organizing center for the different types of EAD dynamics that have been observed. To the best of our knowledge, this work is the first to consider connections between the circadian clock and EADs.

## Methods

### Single-cell model

We used recently published voltage-clamp recordings from guinea pig ventricular myocytes to modify the Sato et al. [57] minimal model of cardiac action potential generation. This conductance-based model describes the dynamics of the membrane potential *V* using the Hodgkin-Huxley modeling formalism and is a three-dimensional system of ordinary differential equations (ODEs):

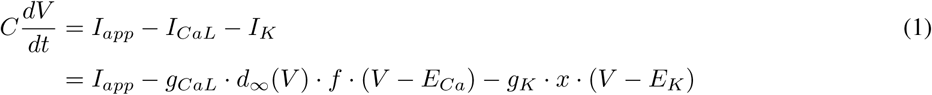

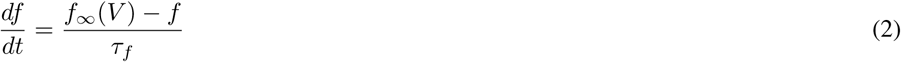

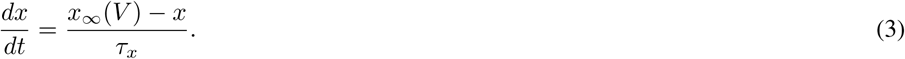

The model includes an inward L-type calcium current (*I*_*CaL*_), an outward potassium current (*I*_*K*_), and an externally applied current *I*_*app*_. Inward sodium current is not included here as it does not impact EAD generation due to the inactivation of this current at depolarized membrane potentials [41]. The calcium current activates instantaneously as a function of voltage, *d*_∞_ (*V*). Inactivation of the calcium current is governed by the gating variable *f* with steady-state inactivation *f*_∞_ (*V*) and time constant *τ*_*f*_. Activation of the potassium current occurs on a slower time scale and is described by the gating variable *x* with steady-state activation *x*_∞_ (*V*) and time constant *τ*_*x*_. The specific membrane capacitance is *C* = 1 *µ*F/cm^2^.

The voltage-dependent activation and inactivation functions are sigmoids given by

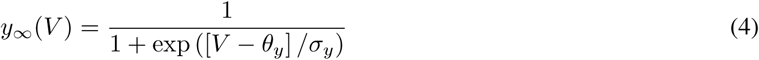

for *y* = *d, f*, and *x*, with half-(in)activation voltages *θ*_*y*_ and slopes proportional to 1/*s*_*y*_.

All of the parameters for the potassium current, except the maximal conductance *g*_*K*_, were kept the same as in the Sato model: *τ*_*x*_ = 300 ms, reversal potential *E*_*K*_ = –80 mV, and activation kinetics *θ*_*x*_ = –40 mV, *σ*_*x*_ = –5 mV. For the calcium current, we set *τ*_*f*_ = 80 ms as in the Sato model, and then fit the remaining parameters to the voltage-clamp data from Chen et al. [9] shown in Figure 1A. In these recordings, Chen et al. measured the L-type calcium current in cardiomyocytes isolated from guinea pigs housed under 12h:12h light:dark cycles, with the lights turned on at Zeitgeber time 0 (ZT0, 7:00 AM) and turned off at ZT12 (7:00 PM). They performed voltage-clamp experiments in the morning (ZT 3) and at night (ZT 15), in which they held cardiomyocytes at -80 mV and then depolarized them in 10 mV increments from -70 to +60 mV. They found that at both times of day, the largest calcium currents were evoked at the +10 mV voltage step. Furthermore, the current density at this voltage step was significantly larger in the morning than at night.

**Figure 1:**
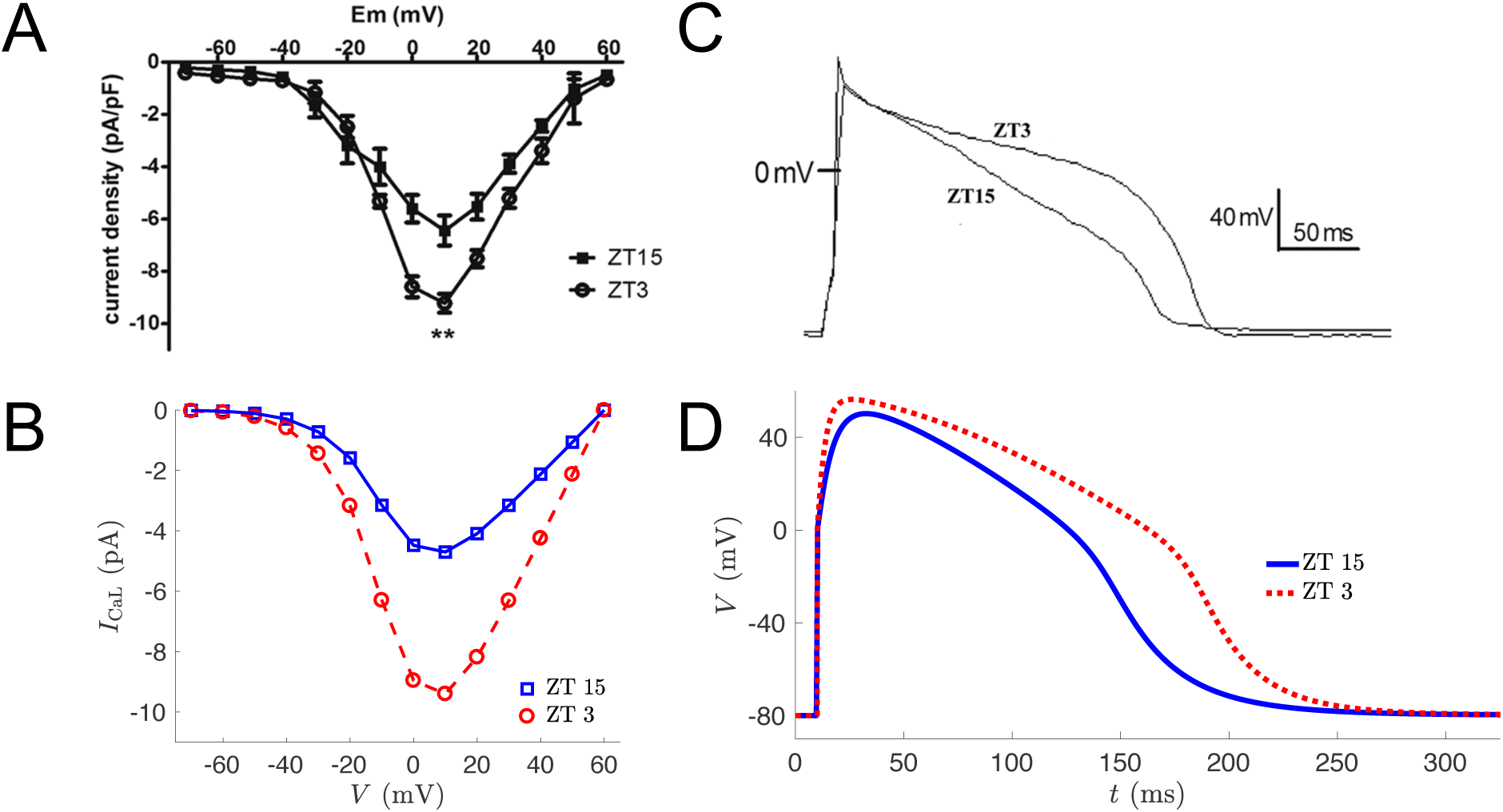
Fitting model parameters to voltage and current-clamp data from guinea pig cardiomyocytes. (***a***) Voltage-clamp data of L-type calcium current from [9]. (***b***) Simulated voltage-clamp experiment with *g*_*CaL*_ = 0.3 mS/cm^2^ (dashed red line and open circles) and *g*_*CaL*_ = 0.15 mS/cm^2^ (solid blue line and open squares) for *g*_*K*_ = 0.1 mS/cm^2^. (***c***) Current-clamp recording of action potentials in guinea pig cardiomyocytes from [9]. (***d***) Simulated current-clamp experiment with *g*_*CaL*_ = 0.3 mS/cm^2^ (dashed red line) and *g*_*CaL*_ = 0.15 mS/cm^2^ (solid blue line) for *g*_*K*_ = 0.1 mS/cm^2^.

We digitized their published I-V curves and normalized the data at ZT 3 and ZT 15 using the peak current density at +10 mV for each time of day. We then averaged these normalized curves to obtain a single curve to use as input to a parameter estimation algorithm. Specifically, we used an unconstrained nonlinear optimization routine (the Nelder-Mead algorithm *fminsearch* in MATLAB) and voltage-clamp simulations to find the parameter values *E*_*Ca*_ = 60 mV, *θ*_*d*_ = –7.3 mV, *σ*_*d*_ = –8.6 mV, *θ*_*f*_ = –13.3 mV, and *σ*_*f*_ = 11.9 mV. These parameter values minimized the squared error between the model-generated *I*_*CaL*_ I-V curve and the average normalized I-V curve from the voltage-clamp data. With the reversal potential and gating kinetics held fixed, we varied the maximal conductance and found that *g*_*CaL*_ = 0.3 mS/cm^2^ produced a model I-V curve (blue curve in Fig. 1B) similar to the experimental data from ZT 3, whereas the *g*_*CaL*_ = 0.15 mS/cm^2^ curve (red curve in Fig. 1B) was similar to the data from ZT 15.

To determine the maximal conductance *g*_*K*_, we performed current-clamp simulations and compared the model traces to the current-clamp recordings from Chen et al. [9] shown in Fig. 1C. For these simulations, we set *I*_*app*_ = 0 and simulated for 10 ms with initial conditions *V*_0_ = –80 mV, *f*_0_ = *f*_∞_ (–80) = 0.9963, and *x*_0_ = *x*_∞_ (–80) = 0.0003354. At *t* = 10 ms, we instantaneously set *V* = 0 to mimic the effect of a stimulating current pulse [34]. We then measure the action potential duration at 90% repolarization (APD90), which is the amount of time it takes for the voltage to return to 90% of its value before the spike. We find that setting *g*_*K*_ = 0.1 mS/cm^2^ with *g*_*CaL*_ = 0.15 mS/cm^2^ (blue curve in Fig. 1D) yields an APD90 for the model that is similar to the APD90 in the experimental data at ZT 15. Furthermore, setting *g*_*K*_ = 0.1 mS/cm^2^ with *g*_*CaL*_ = 0.3 mS/cm^2^ (red curve in Fig. 1D) gives a model APD90 that is very similar to the experimental APD90 at ZT 3.

Simulations of the single-cell model were performed using MATLAB R2017a (The Mathworks Inc., Natick, MA) and *ode15s*, a variable-step, variable-order solver for stiff ODEs.

### Bifurcation analysis

The single-cell model was analyzed by decomposing it into a fast subsystem (Eq. 1–2) and a slow subsystem (Eq. 3), as in [66]. We then treat the slow variable *x* as a bifurcation parameter and study the bifurcation structure of the fast subsystem:

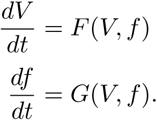

The linearization of this system at a steady state (*V* ^*∗*^, *f* ^*∗*^) is given by the Jacobian matrix

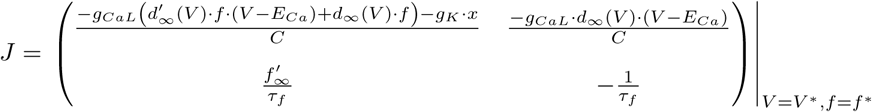

At a steady state, we have that *F* (*V, f*) = *G*(*V, f*) = 0. To find steady states, we set *f* ^*∗*^ = *f* (*V* ^*∗*^) to satisfy *G*(*V* ^*∗*^, *f* ^*∗*^) = 0, and then solve *F* (*V* ^*∗*^, *f* ^*∗*^) = 0 for *V* ^*∗*^:

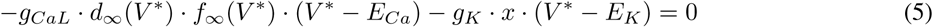

Hopf bifurcation occurs when trace(*J*) = 0 and determinant(*J*) *>* 0. Saddle-node bifurcation occurs when trace(*J*) ≠ 0 and determinant(*J*) = 0. Takens-Bogdanov (TB) bifurcation occurs when Hopf and saddle-node bifurcation points coalesce and the Jacobian matrix has two zero eigenvalues [14]. The conditions for this codimension-2 bifurcation are trace(*J*) = 0 and determinant(*J*) = 0, that is

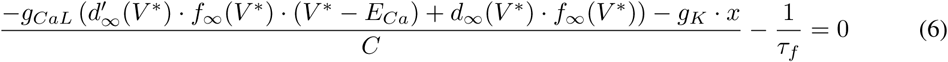

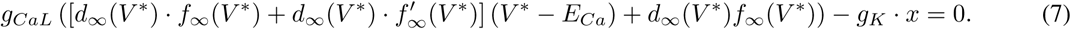

To find TB points, we simultaneously solve Eqs. 5–7. Bifurcations were also identified using the dynamical systems software package XPPAUT [16].

### Circadian variation of maximal conductances

To simulate circadian rhythms in the maximal conductances of the calcium and potassium channels, we assumed a sinusoidal waveform with peak (trough) times of ZT 3 (ZT 15) for calcium and ZT 14 (ZT 2) for potassium:

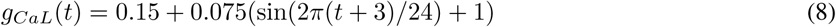

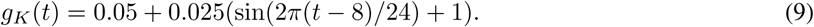

### Spatial model

In cardiac tissue, neighboring cells are electrically coupled through gap junctions. The spatiotemporal evolution of the cellular membrane potential in a 2-D domain can be described by the following reaction-diffusion partial differential equation (PDE):

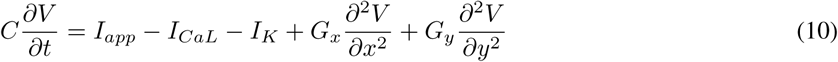

where *G*_*x*_ and *G*_*y*_ are longitudinal and transverse conductances associated with the diffusion terms representing intercellular currents. To simulate a 2-D sheet of guinea pig cardiac tissue, we modified the monodomain reaction-diffusion MATLAB code developed by Hammer [26]. We solved the PDEs numerically on a 128×128 isotropic (*G*_*x*_ = *G*_*y*_ = 25 nS) grid using a finite-difference scheme for spatial derivatives, the explicit Euler method for time derivatives, and Neumann (no-flux) boundary conditions, with a time step of 0.1 ms and a space step of 0.01 cm. Cardiomyocytes are typically 100 *µ*m in length and 10-25 *µ*m in diameter. Thus our simulated tissue size of 1.6384 cm^2^ represents approximately 128 cells in the longitudinal direction and 512 to 1280 cells in the transverse direction (or 65,536 to 163,480 cells in total).

We used an S1-S2 cross-field stimulation protocol, with the first stimulus (S1) delivered to the left boundary of the domain at *t* = 0 with strength *I*_*app*_ = 500 *µ*A/cm^2^ and a duration of 2 ms. The second stimulus (S2), was delivered to the bottom domain boundary at *t* = 810 ms with the same strength as S1 and a duration of 3 ms. This stimulation procedure generates spiral waves in our 2-D domain as shown in Fig. 6.

## Results

### Elevated L-type calcium current in the morning can induce EADs

To investigate the role that circadian rhythmicity of the L-type calcium current plays in the electrical activity of guinea pig cardiomyocytes, we simulated an electrophysiological model of these cells (Eq. 1–3) with maximal conductance values corresponding to morning and evening time points: specifically *g*_*Ca*_ = 0.3 mS/cm^2^ at ZT 3 and *g*_*Ca*_ = 0.15 mS/cm^2^ at ZT 15. We determined these parameter values, along with the gating kinetics of the calcium current (Eq. 4), by fitting voltage-clamp data from Chen et al. [9] as described in the Methods section (see Fig. 1, A and B). This model, with maximal potassium conductance *g*_*K*_ = 0.1 mS/cm^2^, can reproduce the circadian variation in action potential duration observed in current-clamp recordings (see Fig. 1, C and D). In the current-clamp data, APD90 is 11.5% greater at ZT 3 (228.0 ms) than ZT 15 (204.5 ms); in the model, APD90 is 16% greater at ZT 3 (225.5 ms) than ZT 15 (194.4 ms). The model enables us to explore the interaction between the potassium conductance and circadian variation of L-type calcium. We find that if *g*_*K*_ is lowered to 0.05 mS/cm^2^, then the difference between morning and evening becomes more pronounced, with APD90 being 97.5% greater at ZT 3 (579.9 ms) than ZT 15 (293.6 ms), see Fig. 2. Moreover, the action potential at ZT 3 now exhibits secondary voltage depolarizations during the repolarization phase, known as early afterdepolarizations (EADs).

**Figure 2:**
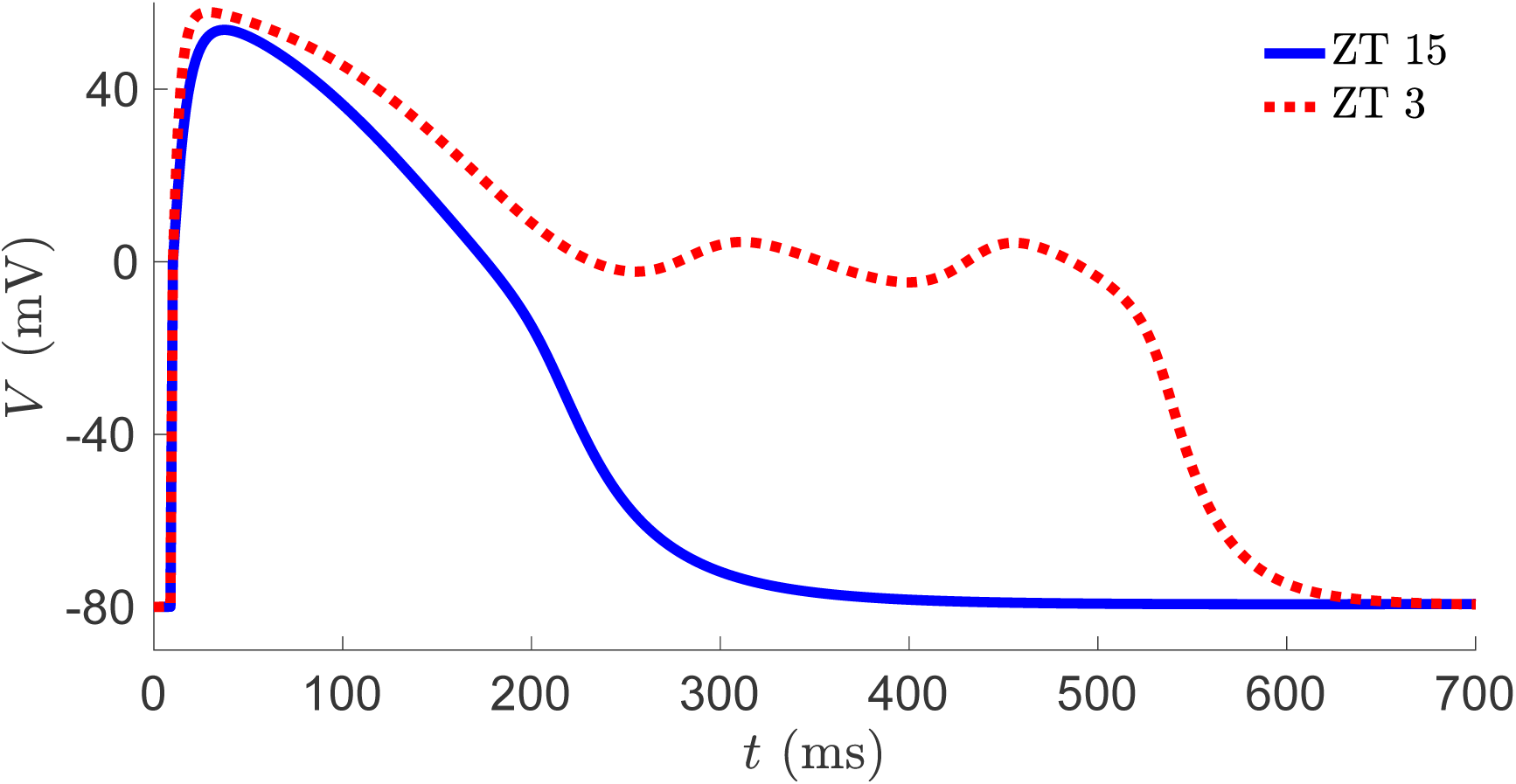
Model exhibits early afterdepolarizations for high *g*_*CaL*_ and low *g*_*K*_. Voltage trajectories from simulated current-clamp experiments with *g*_*CaL*_ = 0.3 mS/cm^2^ (dashed red line) and *g*_*CaL*_ = 0.15 mS/cm^2^ (solid blue line) for *g*_*K*_ = 0.05 mS/cm^2^.

### Dynamics of EAD generation

To understand the dynamical mechanism underlying the generation of these EADs, we follow Tran et al. [66] and perform a fast-slow decomposition of our model. As described in the Methods section, we study bifurcations in the fast (*V, f*) subsystem (Eqs. 1–2) treating the slow variable *x* as a bifurcation parameter. The fast subsystem generally has three fixed points for small values of *x* and one fixed point for large values of *x*, forming a Z-shaped curve in the *V, x* plane (Fig. 3). The curve consists of an upper branch of depolarized fixed points, a middle branch of unstable fixed points, and a lower branch of hyperpolarized stable fixed points. As *x* is increased, the fixed points on the upper branch change from stable (solid red curve) to unstable (dashed black curve) at a subcritical Hopf bifurcation, where unstable limit cycles (open green circles) are born. These unstable periodic solutions are terminated at a homoclinic bifurcation with a saddle point on the middle branch of fixed points. As *x* is increased further, the upper and middle branches of fixed points approach each other and eventually coalesce, destroying these fixed points in a saddle-node bifurcation. With *g*_*CaL*_ = 0.15 mS/cm^2^ and *g*_*K*_ = 0.1 mS/cm^2^, the Hopf and saddle-node bifurcations occur at *x*_*HB*_ = 0.202 and *x*_*SN*_ = 0.275, respectively (3A). When repolarizing, the action potential (AP) trajectory (blue curve) passes through the *V, x* plane to the right of these values (*x > x*_*SN*_), and therefore repolarizes monotonically without EADs. If *g*_*CaL*_ is increased to 0.3 mS/cm^2^ with *g*_*K*_ held fixed (Fig. 3B), the Hopf and saddle-node bifurcation points move to the right (*x*_*HB*_ = 0.376, *x*_*SN*_ = 0.551), but so does the AP trajectory; here the AP repolarizes through the region *x*_*HB*_ *< x < x*_*SN*_ without EADs. Similarly, if *g*_*CaL*_ is held fixed at 0.15 mS/cm^2^ but *g*_*K*_ is reduced to 0.05 mS/cm^2^ (Fig. 3C), the trajectory repolarizes without EADs through the region *x*_*HB*_ = 0.404 *< x < x*_*SN*_ = 0.551. However, if *g*_*CaL*_ is increased to 0.3 mS/cm^2^ and *g*_*K*_ is reduced to 0.05 mS/cm^2^, the model does exhibit EADs. The AP trajectory now repolarizes through a region where *x < x*_*HB*_ = 0.753 and the fast subsystem contains a stable fixed point (Fig. 3D). This fixed point is a stable focus, and the trajectory exhibits damped oscillations until it reaches *x*_*HB*_. To the right of *x*_*HB*_, the fixed point is now an unstable focus and the trajectory exhibits one more voltage peak before repolarizing fully.

**Figure 3:**
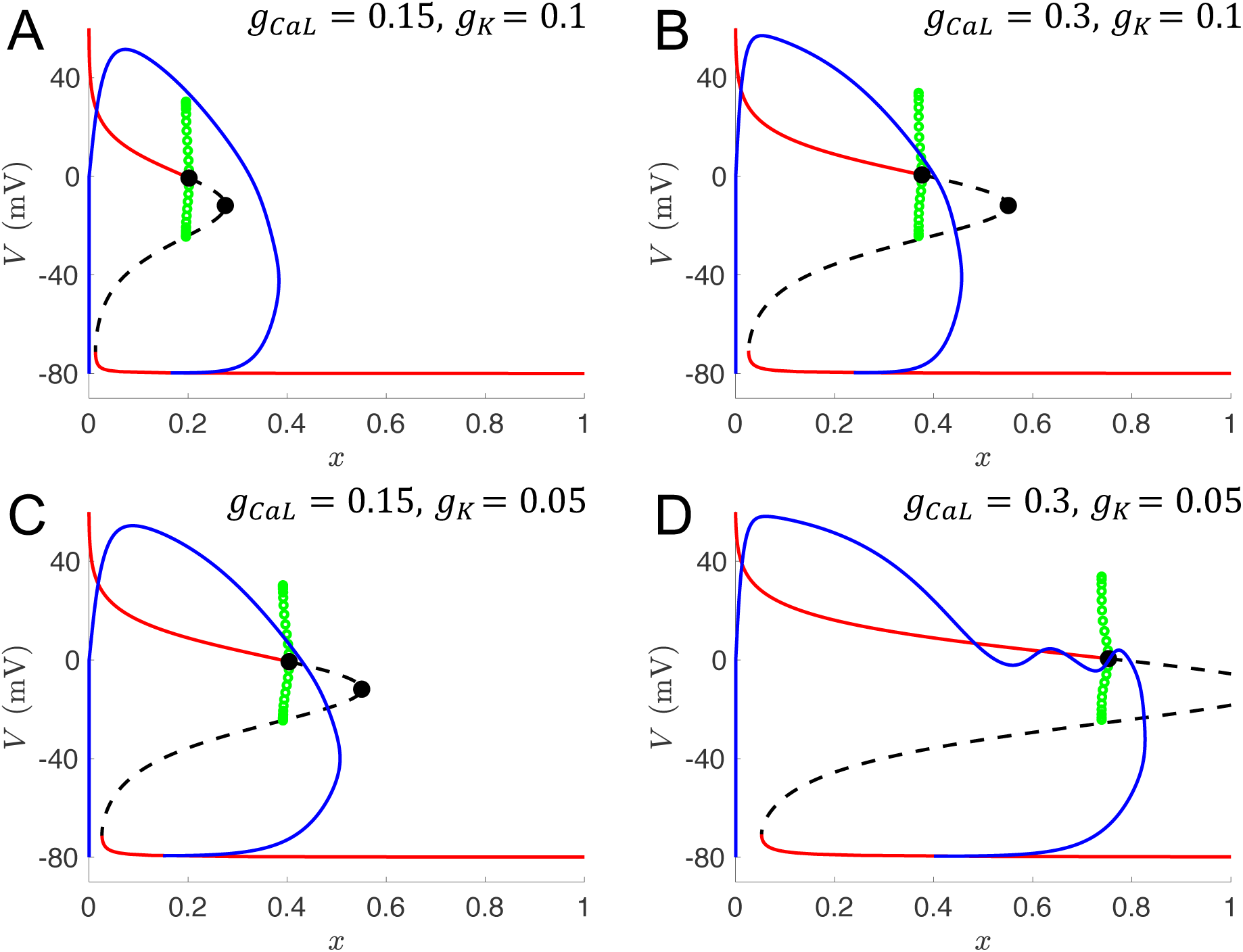
Bifurcation diagrams with bifurcation parameter *x* for various values of *g*_*Ca*_ and *g*_*K*_. Trajectories from the full system (solid blue lines) are projected onto the *x –V* plane and overlayed with steady states of the fast subsystem (solid red lines are stable, dashed black lines are unstable), along with bifurcation points (solid black dots) and unstable periodic orbits (open green circles) emanating from the subcritical Hopf bifurcation. (***a***) Normal APs for *g*_*CaL*_ = 0.15 mS/cm^2^ and *g*_*K*_ = 0.1 mS/cm^2^. (***b***) Increased APD, but not EADs, with increased *g*_*CaL*_. (***c***) Increased APD, but not EADs, with reduced *g*_*K*_. (***d***) EADs with increased *g*_*CaL*_ and reduced *g*_*K*_.

### Takens-Bodganov bifurcation as an organizing center

Several different dynamical mechanisms can give rise to secondary oscillations that grow in amplitude, which is the EAD pattern typically observed in experiments. In the first mechanism to be characterized, stable limit cycles with growing amplitudes emerge from a supercritical Hopf bifurcation in the fast subsystem [66]. More recently, Kügler [34] demonstrated that EADs with growing amplitudes can also arise either from a delayed subcritical Hopf bifurcation in the fast subsystem, or along the unstable manifold of a saddle-focus fixed point in the full system. In the latter case, there is no Hopf bifurcation. The EADs explored earlier in this paper are of the subcritical Hopf type; see the bifurcation diagrams in Fig. 3. In these diagrams, the Hopf bifurcations occur relatively near a saddle-node bifurcation. This suggests that by varying another parameter in conjunction with the bifurcation parameter *x*, the Hopf and saddle-node bifurcations can be made to coalesce in a Takens-Bogdanov (TB) bifurcation. Indeed, Fig. 4A shows that the Hopf and saddle-node bifurcation points approach and collide with each other as *x* and *g*_*CaL*_ are decreased simultaneously, with the TB bifurcation occurring at *x* = 0.411, *g*_*CaL*_ = 0.0224 mS/cm^2^. We simulated the model with parameters chosen near the TB bifurcation point (*g*_*CaL*_ = 0.02, *g*_*K*_ = 0.01, *θ*_*x*_ = –40, *τ*_*x*_ = 1100) and observed EADs as shown in Fig. 4B-C. The eigenvalues of the full system linearized at the fixed point (*V* ^*∗*^, *f* ^*∗*^, *x*^*∗*^) = (–12.74, 0.4882, 0.3664), which corresponds to a location near the saddle-node bifurcation point shown in Fig. 4B, are *λ*_1,2_ = 0.0025 ±0.0066*i* and *λ*_3_ = –0.0068. Thus, this fixed point is classified as a spiral saddle of index 2 [27]. The EADs arise due to the spiraling movement of the trajectory caused by the unstable manifold of the saddle focus, which is the second EAD-generating mechanism found by Kügler [34]. In this way, knowledge of the TB bifurcation can help identify parameter sets that produce different types of EAD dynamics.

**Figure 4:**
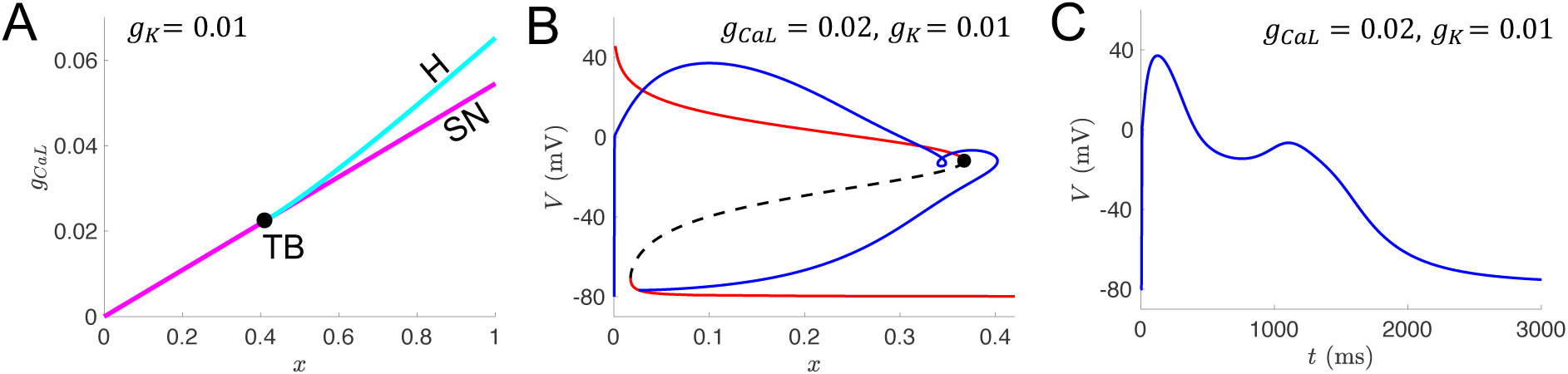
EAD generation via a different dynamical mechanism near Takens-Bogdanov bifurcation point. (***a***) Two-parameter bifurcation diagram showing the location of Hopf bifurcations (H, cyan curve) and saddle-node bifurcations (SN, magenta curve) for bifurcation parameters *g*_*CaL*_ and *x*, with *g*_*K*_ = 0.01 mS/cm^2^. The Hopf and SN curves coalesce at a codimension-2 Takens-Bogdanov bifurcation (TB, solid black dot). (***b***) Trajectory exhibiting an EAD (solid blue line) from the full system projected onto the *x –V* plane and overlayed with steady states of the fast subsystem (solid red lines are stable, dashed black lines are unstable), along with the saddle-node bifurcation point (solid black dot), for maximal conductance parameters (*g*_*CaL*_ = 0.02 mS/cm^2^, *g*_*K*_ =0.01 mS/cm^2^) chosen near the TB bifurcation point shown in panel (a). (***c***) Voltage time course of the EAD trajectory shown in panel (b).

### Circadian variation of calcium and potassium currents

The voltage-clamp experiments of Chen et al. [9] revealed a day/night difference in L-type calcium current, with larger currents in the morning (ZT 3) than at night (ZT 15). Correspondingly, they observed longer duration action potentials in their current-clamp recordings at ZT 3 than at ZT 15. We used our model to simulate action potentials across the circadian cycle by assuming that *g*_*CaL*_ follows a sinusoidal waveform with a period of 24 hours (Eq. 8), with a maximum of *g*_*CaL*_ = 0.3 at ZT 3 and a minimum of *g*_*CaL*_ = 0.15 at ZT 15. With *g*_*K*_ held fixed at 0.1, we find that EADs do not occur at any time of day. Figures 5A and 4B show the APD90 values across a circadian cycle with *g*_*K*_ = 0.1 and *g*_*K*_ = 0.05 respectively. With *g*_*K*_ = 0.05, we find that EADs occur over a large portion of the day (∼8 hours), specifically from ZT 23 to ZT 7. We simulated with two different *g*_*K*_ values to represent the heterogeneity in potassium channel expression that has been found across different cells of the ventricular myocardium [69] or among different guinea pigs. However, it is plausible that *g*_*K*_ may also be under circadian control, since it has been shown that IKs is modulated by K^+^ channel interacting protein 2 (KChIP2) and that KChIP2 expression follows a circadian rhythm in mouse ventricles. In particular, co-expression of KChIP2 reduces IKs [39], and KChIP2 expression is higher at night than in the morning [29]. Thus we modeled a circadian rhythm in potassium channel conductance by assuming *g*_*K*_ follows a sinusoidal waveform with a period of 24 hours (Eq. 9), with a maximum of *g*_*K*_ = 0.1 at ZT 14 and a minimum of *g*_*K*_ = 0.05 at ZT 2. Figure 5C shows the APD90 values across a circadian cycle with rhythms in both *g*_*CaL*_ and *g*_*K*_. We find that EADs occur over an *∼*5-hour portion of the day, from ZT 1 to ZT 6.

**Figure 5:**
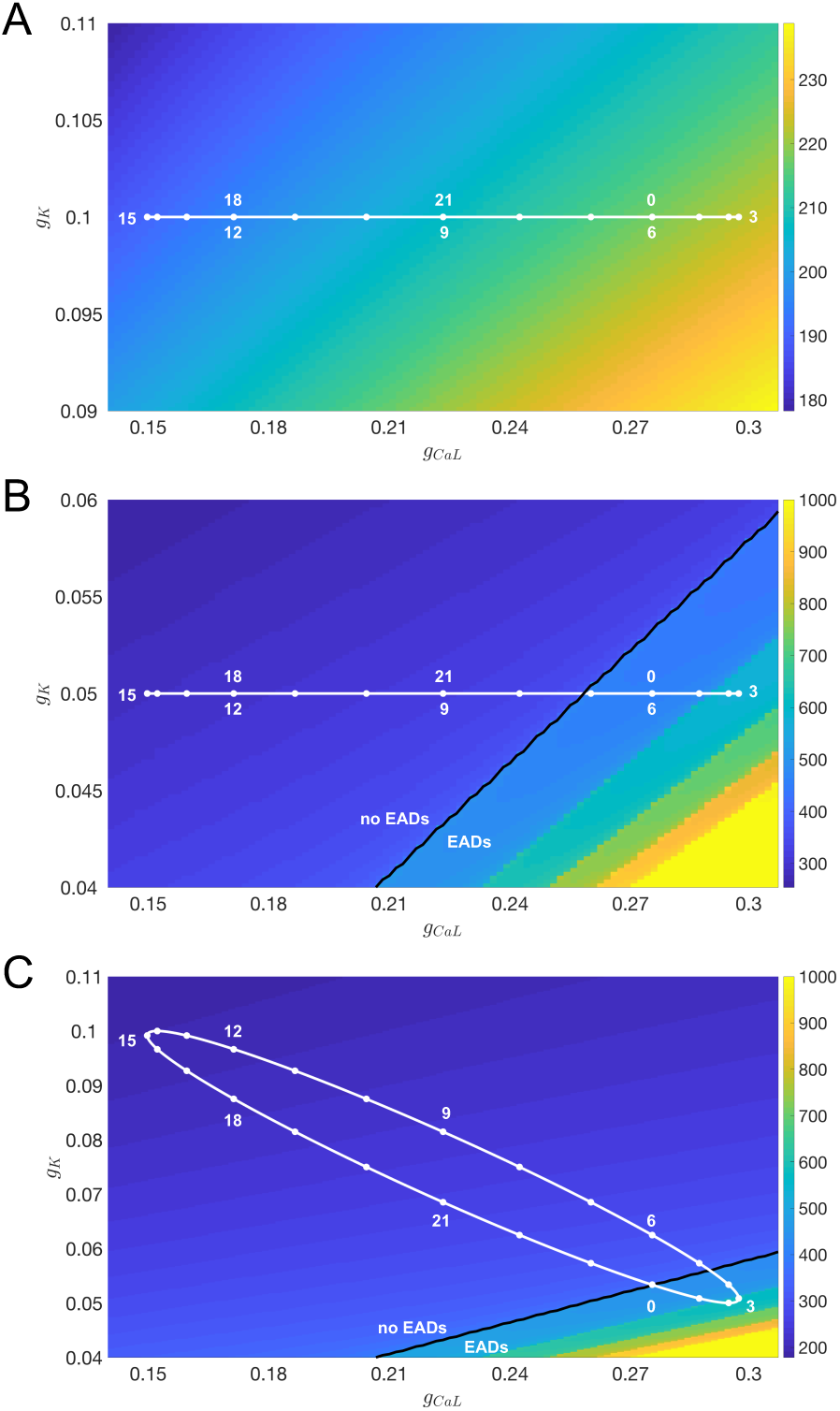
Action potential durations and early afterdepolarizations over a circadian cycle. Colorbar indicates APD (in ms), solid white dots are hourly ZT markers, and black lines separate regions of parameter space with and without EADs. (***a***) Circadian variation of *g*_*CaL*_ (Eq. 8) with *g*_*K*_ = 0.1 mS/cm^2^ does not result in EADs. (***b***) Circadian variation of *g*_*CaL*_ (Eq. 9) with reduced *g*_*K*_ = 0.05 mS/cm^2^. EADs occur between ZT 23 and ZT 7. (***c***) Circadian variation of both *g*_*CaL*_ and *g*_*K*_ (Eqs. 8-9). EADs occur between ZT 0 and ZT 5.

**Figure 6:**
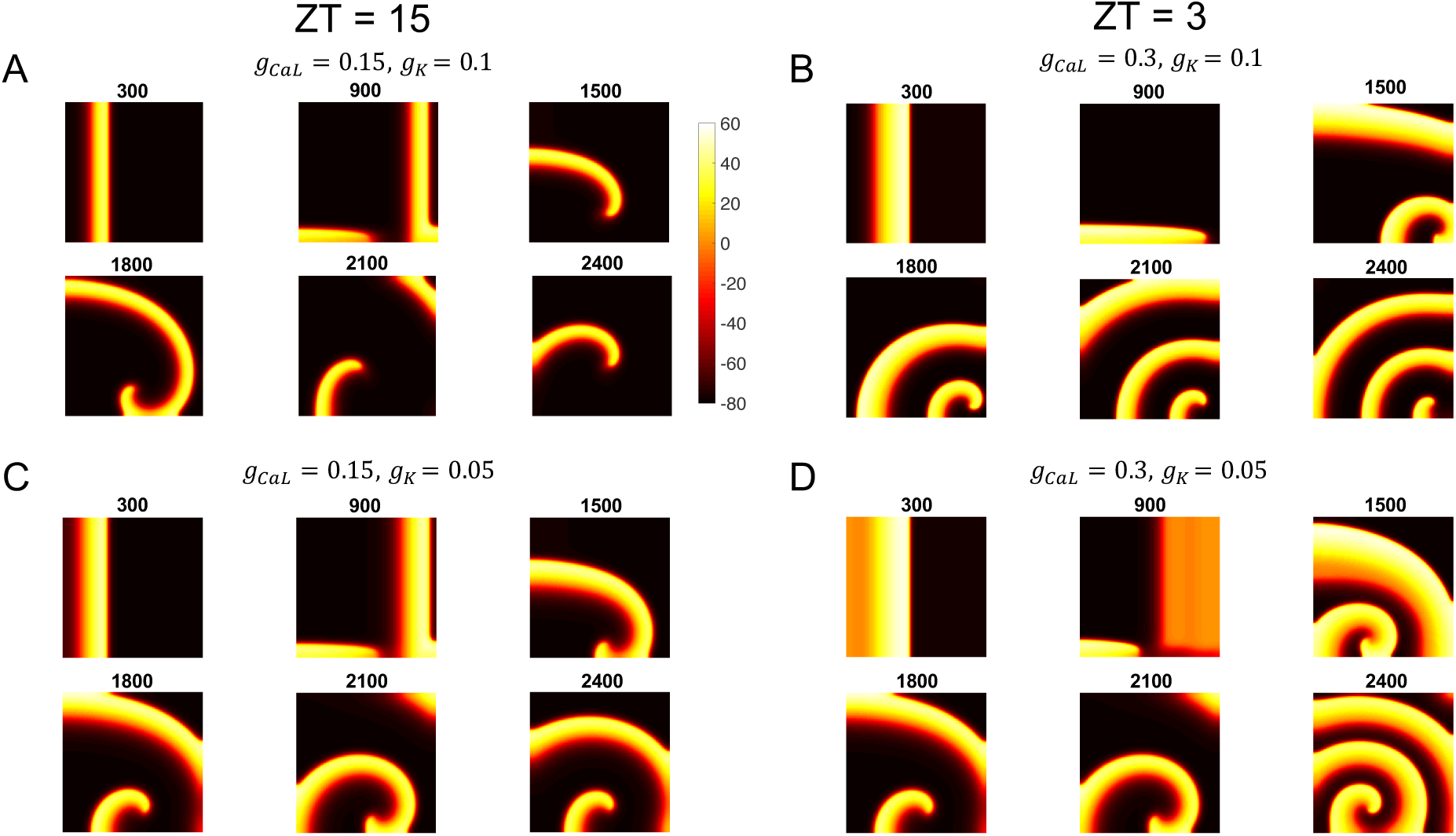
Spiral waves in a 2-D domain. Colorbar indicates membrane voltage (mV) at snapshots of *t* = 300, 900, 1200, 1500, 1800, 2100, and 2400 ms for simulations of Eq. 10 on a 128 × 128 grid under an S1-S2 cross-field stimulation protocol. (***a***) Parameters corresponding to ZT 15 (*g*_*CaL*_ = 0.1 mS/cm^2^) with *g*_*K*_ = 0.1 mS/cm^2^. (***b***) Parameters corresponding to ZT 3 (*g*_*CaL*_ = 0.3 mS/cm^2^) with *g*_*K*_ = 0.1 mS/cm^2^. (***c***) Parameters corresponding to ZT 15 (*g*_*CaL*_ = 0.1 mS/cm^2^) with reduced *g*_*K*_ = 0.05 mS/cm^2^. (***d***) Parameters corresponding to ZT 3 (*g*_*CaL*_ = 0.3 mS/cm^2^) with reduced *g*_*K*_ = 0.05 mS/cm^2^, which produce EADs in the isolated single-cell model as shown in Fig. 3D.

### EADs lead to pathological electrical activity in 2-D tissue simulations

To explore whether the single-cell EADs triggered by circadian variation of ion channel conductances leads to abnormal electrical activity in cardiac tissue, we simulated a 2-D spatial domain as described in the Methods section. An S1-S2 stimulation protocol triggered spiral waves at both ZT 3 (*g*_*CaL*_ = 0.3 mS/cm^2^) and ZT 15 (*g*_*CaL*_ = 0.15 mS/cm^2^) with either normal (*g*_*K*_ = 0.1 mS/cm^2^) or low (*g*_*K*_ = 0.05 mS/cm^2^) potassium conductance (Fig. 6). Of these 4 scenarios, only the ZT 3 low *g*_*K*_ combination exhibited EADs in the spatial model (Fig. 7). In addition, this combination led to the steepest APD restitution curve (Fig. 8), a commonly used indicator of the propensity for ventricular tachyarrhythmias [22, 30, 49]. To test this propensity, we next simulated heterogeneity in potassium channel conductance across the tissue with the leftmost 80% of the domain set to *g*_*K*_ = 0.05 mS/cm^2^ and the rightmost 20% set to *g*_*K*_ = 0.1 mS/cm^2^.

**Figure 7:**
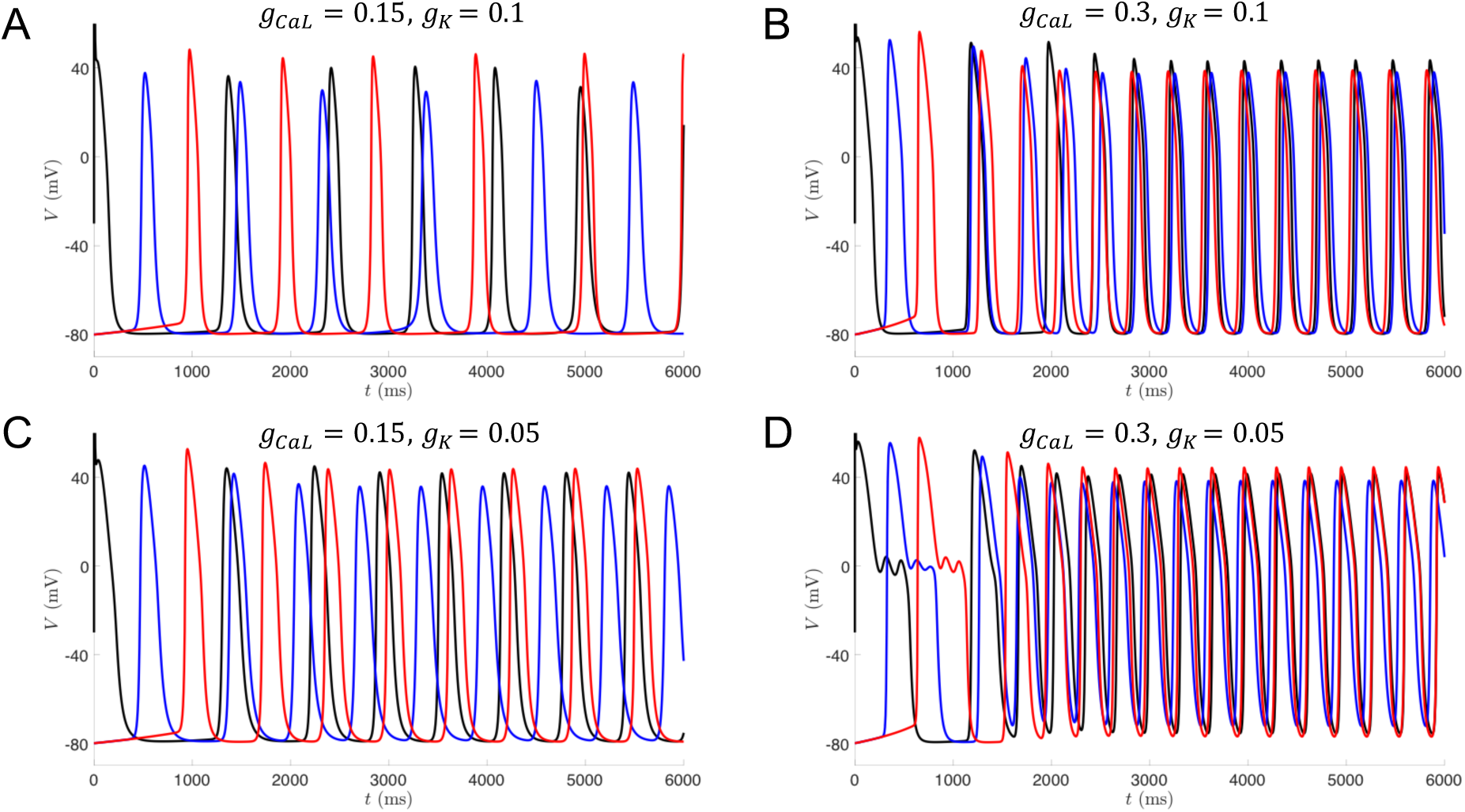
Voltage trajectories for 3 locations in the 2-D spatial model. Leftmost (black), center (blue), and rightmost (red) grid points for the middle row of the 128 × 128 domain shown in Fig. 6. (***a***) Parameters corresponding to ZT 15 (*g*_*CaL*_ = 0.1 mS/cm^2^) with *g*_*K*_ = 0.1 mS/cm^2^. (***b***) Parameters corresponding to ZT 3 (*g*_*CaL*_ = 0.3 mS/cm^2^) with *g*_*K*_ = 0.1 mS/cm^2^. (***c***) Parameters corresponding to ZT 15 (*g*_*CaL*_ = 0.1 mS/cm^2^) with reduced *g*_*K*_ = 0.05 mS/cm^2^. (***d***) Parameters corresponding to ZT 3 (*g*_*CaL*_ = 0.3 mS/cm^2^) with reduced *g*_*K*_ = 0.05 mS/cm^2^.

**Figure 8:**
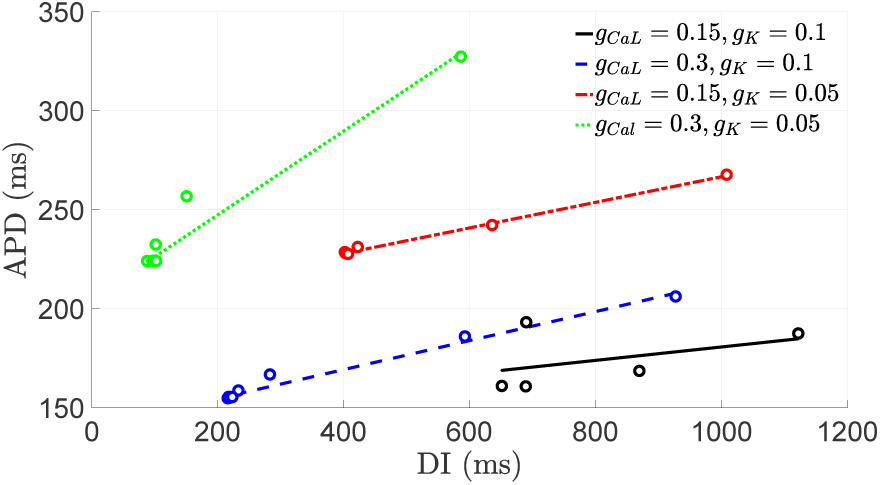
APD restitution curves from the 2-D spatial model. APD and Diastolic Interval (DI) was calculated for the leftmost (black) voltage trajectories shown in Fig. 7. Open circles denote (DI,APD) values from each of the four simulations. Linear fits to the data points for the simulations shown in Fig. 7A (solid black), 7B (dashed blue), 7C (dashed-dotted red), and 7D (dotted green).

At ZT 15, the solution consists of a single spiral wave (Fig. 9A). However, at ZT 3, multiple spiral waves are born and collide into each other (Fig. 9B). This type of spiral wave break-up has been associated with ventricular fibrillation.

**Figure 9:**
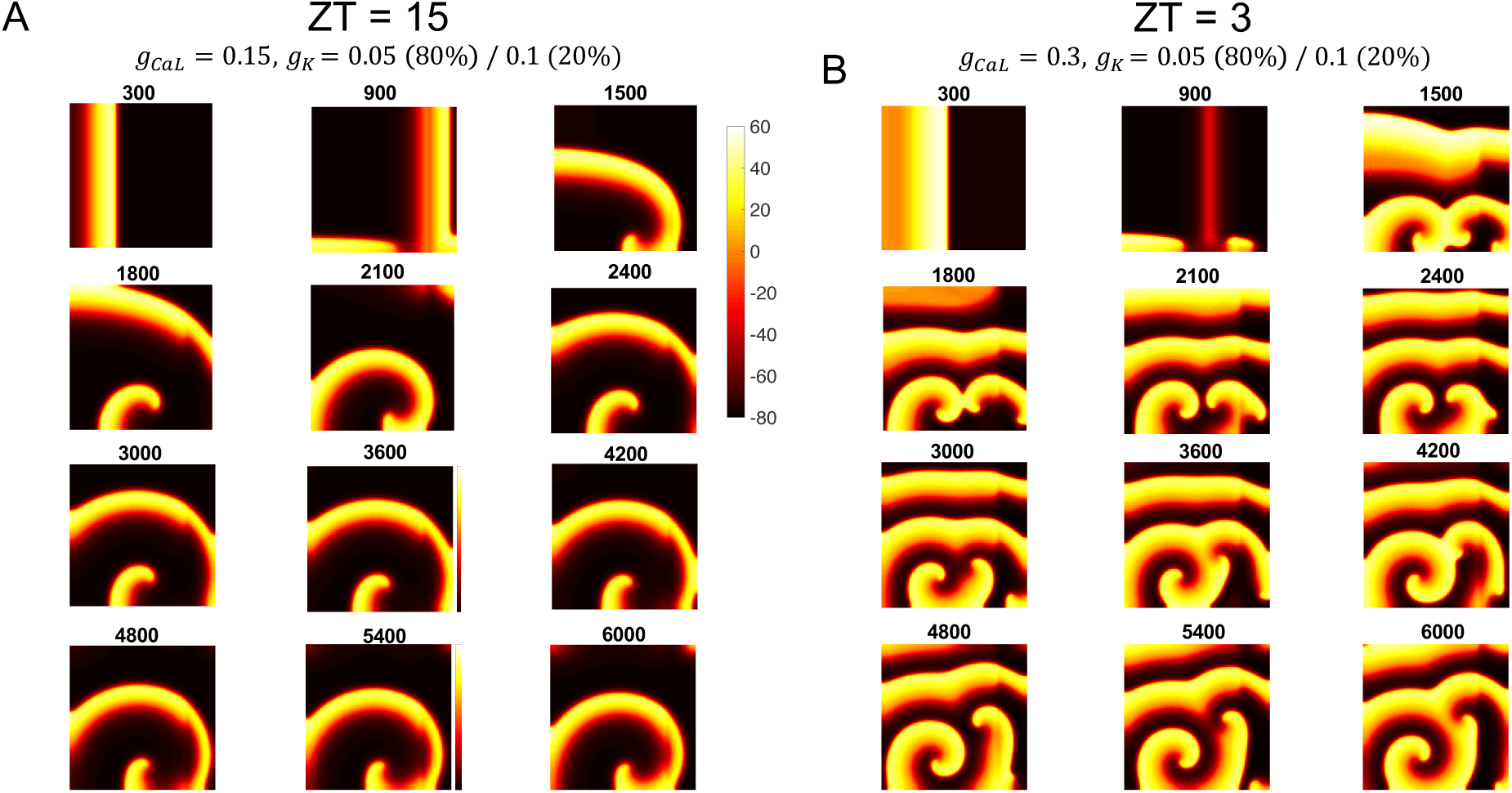
Spiral wave break-up in a 2-D domain. Same simulation and stimulation protocol as Fig. 6 but with heterogeneity in potassium conductance across the domain; *g*_*K*_ = 0.05 mS/cm^2^ for the leftmost 80% of the domain and *g*_*K*_ = 0.1 mS/cm^2^ for the rightmost 20%. (***a***) Single spiral wave for parameters corresponding to ZT 15 (*g*_*CaL*_ = 0.15 mS/cm^2^). (***b***) Break-up into multiple spiral waves for parameters corresponding to ZT 3 (*g*_*CaL*_ = 0.3 mS/cm^2^).

## Discussion

### Cardiac arrhythmogenesis and circadian rhythms

Epidemiological studies have shown that the occurrence of life-threatening cardiac arrhythmias, such as ventricular tachycardia and ventricular fibrillation, exhibits time-of-day dependence with a peak in the morning hours [52]. For example, episodes of ventricular tachyarrhythmias recorded in patients with implantable cardioverter defibrillators were significantly increased between 8:00 and 11:00 AM [32]. Controlled laboratory studies indicate that the time-of-day fluctuations in adverse cardiovascular events are not solely due to daily rhythms in behavior and the external environment, suggesting that internally generated circadian oscillations influence cardiac arrhythmogenesis [8]. Normal electrical properties of the heart, such as electrocardiogram waveforms and heart rate, also demonstrate robust circadian rhythms [15, 13]. The circadian system could exert this influence through two primary mechanisms: (1) regulation of cardiac electrophysiology by the central circadian clock in the hypothalamus through neurohumoral factors and the autonomic nervous system, or (2) local circadian clocks in cardiomyocytes themselves driving circadian rhythms in ion channel expression [3]. In this paper we considered the latter mechanism and explored how circadian rhythms in calcium and potassium conductances affect ventricular myocyte electrical activity across the day/night cycle.

### Local cardiac circadian clock

Circadian clocks have been found in mammalian tissues throughout the body, including the heart. These peripheral clocks operate using the same molecular machinery as the central clock in the suprachiasmatic nucleus (SCN). The basic mechanism is a negative feedback loop in which the protein products of the clock genes *Per* and *Cry* inhibit their own production by repressing their transcriptional activator complex CLOCK-BMAL1. The timescales of the biochemical processes involved in this transcription-translation feedback loop lead to oscillations in the abundance of PER and CRY proteins with a period of approximately 24 hours. The expression of many other genes and proteins that are not necessarily integral to the clock mechanism are also influenced by CLOCK-BMAL1 and exhibit ∼24-hour oscillations. Such clock-controlled genes (CCGs), including those that encode ion channels, can then modulate cellular processes in a time-of-day-dependent manner [76]. Oscillations in the expression of core circadian clock genes have been observed in the intact heart, cultured myocardial tissue, and isolated cardiomyocytes [3]. Studies in mice with cardiomyocyte-specific CLOCK mutations (CCM) and BMAL1 knockouts (CBK) demonstrate that 10% of the cardiac transcriptome is regulated by local circadian clocks in the heart [4, 75]. Through these CCGs, the cardiomyocyte circadian clock impacts a variety of key cellular functions, including cardiac metabolism, signal transduction, contractility, and electrophysiology [45].

### Circadian transcription of cardiac ion channels

Several ion channel subunits exhibit circadian rhythms in expression within the ventricles of animal models [3]. The levels of transcripts associated with Na^+^ current (*Scna5*, Nav1.5, *I*_*Na*_) [59], L-type Ca^2+^ current (*Cacna1c* and *Cacna1d*, Cav1.2 and Cav1.3, *I*_*CaL*_) [9, 4], transient outward K^+^ current (*Kcnd2*, Kv4.2, *I*_*to*_) [65], ultra-rapidly activating delayed rectifier K^+^ current (*Kcna5*, Kv1.5, *I*_*Kur*_) [74], rapidly activating delayed rectifier K^+^ current (*Kcnh2*, Kv11.1, *I*_*Kr*_) [58], two-pore background K^+^ current (*Kcnk3*, K2p3.1, *I*_*K*2*p*_) [65], and gap junction current (*Gja5* and *Gja1*, connexins Cx40 and Cx43) [64] oscillate over a 24-hour period. In some cases, rhythms in channel subunit gene expression have been related to day/night differences in electrophysiological properties and cardiac pacemaking. For example, elevated Kcna5 and Kcnd2 protein levels at ZT6 and ZT18, respectively, correlate with increased steady-state currents for *I*_*to*_ and *I*_*Kur*_ at those time points [74]. Potassium Channel Interacting Protein 2 (KChIP2), a regulator of *I*_*to*_, has been implicated in the circadian rhythm of cardiac repolarization. Jeyaraj et al. [29] showed that Kruppel-like factor 15 (Klf15) is a CCG that directly regulates KChIP2 expression, and that deletion of Klf15 abolishes the circadian rhythm in QT interval and increases suspectibility of mice to ventricular arrhythmias. However, Gottlieb et al. [23] concluded that KChIP2 is not a mechanistic link between the cardiac circadian clock and ventricular repolarization and arrhythmogenesis, based on their finding that KChIP2-deficient mice still have a circadian rhythm in QT interval. Rather, they suggest that Klf15 expression controls the transcription of other genes responsible for the circadian rhythm in repolarization and susceptibility to arrhythmias.

### Circadian variation of L-type Ca2+ current

In this paper we focused on circadian regulation of L-type Ca^2+^ channels, due to the evidence supporting local cardiac clock control of these channels and the importance of L-type current for cardiac pacemaking. The *α*1D subunit of the L-type channel (Cacna1d) shows circadian variation in both mRNA and protein expression levels in the hearts of wild-type mice, but not in the hearts of CCM mice [4]. In guinea pigs, the *α*1C subunit of the L-type channel (Cacna1c) is rhythmic at the protein level with a peak at ZT3, which correlates to larger L-type calcium current at that time point [9]. Voltage-gated L-type Ca^2+^ channels have also been proposed as a link between circadian oscillations in electrical activity and the molecular clock in SCN neurons [51, 48, 11, 2] and retinal photoreceptors [31].

Although circadian variation of potassium channel expression has been observed in mouse and rat ventricles, the voltage-clamp studies of Chen et al. [9] did not find a significant time-of-day dependence for the major outward potassium currents (I_Ks_ and I_Kr_) in guinea-pig ventricular myocytes. Thus, in most of this paper we assume the potassium current to be constant throughout the day/night cycle. Instead, we consider the effect of circadian variation in

I_CaL_ in the presence of lower or higher levels of I_Ks_, reflecting the heterogeneity in potassium channel expression one might expect to find across different cells or individuals. Nonetheless, in one set of simulations we do explore the effect of antiphase circadian variation of I_CaL_ and I_Ks_ (see Fig. 5).

### Mathematical analysis of EADs

Mathematical modeling studies have shown that increased inward calcium current and decreased outward potassium current can elongate the cardiac AP and produce the pathological voltage oscillations known as early afterdpolarizations (EADs) [53, 68, 37, 28]. To understand the genesis of EADs, minimal models of the cardiac AP have been analyzed using dynamical systems tools such as slow-fast decomposition and bifurcation theory. Tran et al. [66] showed that EADs involve supercritical Hopf and homoclinic bifurcations in the fast subsystem, and claimed that under periodic pacing the homoclinic bifurcation leads to chaotic behavior. Sato et al. [57] argued that deterministic chaos, rather than random fluctuations due to noise, is the primary cause of the irregular EAD dynamics frequently seen in cardiac experiments [57]. Kügler [34] showed that EADs can also arise from alternative dynamical mechanisms, such as delayed subcritical Hopf or limit point bifurcations of the fast subsystem. Furthermore, Kügler et al. [35] argued that a cascade of period doubling bifurcations underlies EAD chaos in both periodically paced and unpaced cardiomyocytes. These studies all decomposed the full model into fast and slow subsystems with a single gating variable in the slow subsystem. Kügler et al. [36] performed a slow-fast decomposition with two slow variables, and proposed a folded-node singularity of the slow flow as a novel mechanism for EAD generation. Vo and Bertram [70] analyzed the same model treating two variables as slow and also attributed EADs to folded-node singularities and their associated canard orbits. They demonstrated that the appearance of dynamical chaos under periodic pacing can be understood using the theory of candard-induced mixed-mode oscillations [5].

In this paper, we utilized the same 3-variable model for cardiac APs introduced in [57] and studied in [34, 70], but we refit the parameters of the L-type calcium current to the voltage-clamp data of [9]. With these parameters, when the model is analyzed with a 1-slow-2-fast structure, the EADs are generated by a subcritical Hopf bifurcation in the fast subsystem. This is one of the EAD mechanisms explored in [34]. We then showed that a Takens-Bogdanov (TB) bifurcation is present in this model, and that near the TB point we can find EADs generated by the unstable manifold of a saddle-focus fixed point of the full system. This is the other EAD mechanism explored in [34]. Thus, the TB bifurcation serves as an organizing center for the dynamics and helps connect some of the different types of EADs that have been observed previously.

### Modeling of cardiac tissue

To study how circadian variation of ionic conductances affects cardiac excitability at the tissue level, we simulated a 2-D spatial model using reaction-diffusion PDEs and an S1-S2 stimulation protocol. The spatial model exhibited spiral wave solutions at both circadian time points (ZT3 and ZT15) and with both low and high potassium conductance (*g*_*K*_ = 0.05 and 0.1). Under the conditions where the single-cell model exhibits EADs (ZT3 with *g*_*K*_ = 0.05), the spiral waves in the spatial model had a faster propagation speed, analogous to the heart beating faster as during ventricular tachycardia. When spatial heterogeneity in potassium conductance was introduced, the time of day where the single-cell model exhibits EADs produced spiral wave break-up, a behavior associated with ventricular fibrillation [17]. It is generally accepted that EADs at the cellular level can lead to arrhythmias, such as polymorphic ventricular tachycardias (PVT) and Torsade de Pointes (TdP), at the tissue level [70]. Modeling studies have shown that single-cell EADs can cause wave initiation, that these EADs can synchronize to form 2D wave patterns, and that meandering waves in heterogeneous tissue can give rise to the classic ECG appearances of PVT and TdP [71, 12, 7]. Vandersickel et al. performed a systematic study of single-cell EAD excitations and their 2D manifestations in a model of human ventricular tissue. However, there are still many open questions about how EADs progress to perpetuating arrhythmias [67].

### Conclusion and future directions

The main finding of this paper is that circadian rhythms in L-type calcium conductance can lead to early afterdepolarizations at certain times of the day in a model of guinea pig ventricular myocytes. To our knowledge, this is the first study to consider how the cardiomyocyte circadian clock influences the genesis of EADs. We propose that circadian rhythms in EAD occurrence may contribute to the time-of-day-dependent patterns observed in ventricular tachyarrhythmias and sudden cardiac death. However, to establish this connection there are some limitations of our study that would need to be addressed, as discussed below.

First, Zeitgeber times (ZT) for the guinea pig experiments need to be related to real-world time for humans. Guinea pigs are a commonly used animal model for cardiac electrophysiology, as the shape of guinea pig action potentials are more similar to human APs than are the APs of smaller rodents such as mice. On the other hand, guinea pigs are not a commonly used animal model for circadian experiments, and they don’t have particularly strong sleep/wake rhythms [10]. Guinea pigs are crepuscular, meaning they are most active at dawn and dusk and are neither nocturnal nor diurnal [38]. To extrapolate results from studies with nocturnal animal species to diurnal humans, a phase shift of approximately 12 hours is often assumed. For example, in a study assessing the relevance of circadian rhythms for administration of chemotherapy, 7 hours after light onset (ZT7) for mice was taken to be “middle of the night” for humans [24]. However, a recent study found that many cardiovascular drug targets exhibit circadian rhythms in gene expression with a similar phase relationship in mouse and human heart tissue, including several genes encoding L-type Ca^2+^ channel subunits [55]. In our guinea pig simulations, EADs occurred between ZT0 and ZT5 (Fig. 5C). Assuming a similar phase relationship between guinea pigs and humans, this corresponds to an increased likelihood of EAD-induced arrhythmias in the first few hours after waking up, in accordance with the peak time window for sudden cardiac death found in epidemiological studies [47, 72, 63].

Second, in this study we employed a minimal model of cardiac AP generation consisting of a single Ca^2+^ current and a single K^+^ current. The advantage of this approach is that the low dimensionality of the model facilitates bifurcation analysis and an understanding of how circadian rhythms affect the dynamics underlying EAD generation. A disadvantage is that the model is lacking descriptions of some specific types of ion channels that may be relevant for daily variation of cardiac electrical properties. For example, Nav1.5 sodium channels and Kv11.1 (mERG) potassium channels display circadian rhythms in transcription in mouse hearts [59, 58]. Moreover, cardiomyocyte-specific deletion of *Bmal1* abolishes circadian oscillations in both of these channels, suggesting they are under the control of the local cardiac molecular clock [3]. In future work, we plan to investigate how circadian rhythms in these conductances affect the propensity for EADs and arrhythmias in more detailed models of cardiac electrophysiology that include many types of ionic currents and intracellular calcium dynamics.

Due to the critical role that I_CaL_ plays in EAD formation, L-type Ca^2+^ channels have been identified as a promising therapeutic target for suppressing EADs and their arrhythmogenic consequences [43, 42, 44]. Based on the results of our study, we suggest that special attention should be paid to the time of day that drugs targeting L-type channels for EAD suppression are taken in order to enhance their effectiveness. Tailoring the timing of drug administration based on circadian factors, known as chronomedicine or chronopharmacology, is an emerging area of precision medicine with clinical trials showing dosing-time-dependent efficacy or toxicity across several conditions, including hypertension and other cardiovascular disorders [54, 6]. Computational modeling of how the circadian clock modulates therapeutic targets can be used to help predict the optimal dosage time to maximize efficacy or minimize side effects [1].

Cardiotoxicity is the leading cause of drug development discontinuation and withdrawal of drugs from the market [54, 18]. Multiple drugs that have been pulled from the market for causing fatal TdP have the unintended side effect of blocking Kv11.1 (hERG) potassium channels, and screening for ERG block is now a mandatory requirement for new pharmaceuticals [46]. ERG block is a sensitive but not specific measure of TdP risk, i.e. it gives few false negatives but false positives may be preventing safe drugs from entering the market [50]. The Comprehensive *in vitro* Pro-arrhythmia Assay (CiPA) is a new global initiative to create guidelines for the assessment of drug-induced TdP that recommends a central role for computational modeling of ion channels and *in silico* evaluation of compounds [19, 40]. As noted above, many cardiac ion channels exhibit circadian oscillations, including ERG. Thus, we propose that circadian clock modeling should be incorporated into the CiPA paradigm for assessing drug-induced cardiotoxicity.

## Acknowledgments

This material is based upon work supported by the National Science Foundation under Grant No. DMS 1555237 and the U.S. Army Research Office under Grant No. W911NF-16-1-0584. C.O.D. gratefully acknowledges the financial support of the US-UK Fulbright Commission and of the EPSRC via grant EP/N014391/1.

## References

[1] A. Ballesta, P. F. Innominato, R. Dallmann, D. A. Rand, and F. A. Lévi. Systems chronotherapeutics. Pharmacological Reviews, 69(2):161–199, 2017.

[2] M. Belle and C. Diekman. Neuronal oscillations on an ultra-slow timescale: Daily rhythms in electrical activity and gene expression in the mammalian master circadian clockwork. European Journal of Neuroscience, 48:2696–2717, 2018.

[3] N. Black, A. D’Souza, Y. Wang, H. Piggins, H. Dobrzynski, G. Morris, and M. R. Boyett. Circadian rhythm of cardiac electrophysiology, arrhythmogenesis, and the underlying mechanisms. Heart Rhythm, 16(2):298–307, 2019.

[4] M. S. Bray, C. A. Shaw, M. W. Moore, R. A. Garcia, M. M. Zanquetta, D. J. Durgan, W. J. Jeong, J. Y. Tsai, H. Bugger, D. Zhang, A. Rohrwasser, J. H. Rennison, J. R. Dyck, S. E. Litwin, P. E. Hardin, C. W. Chow, M. P. Chandler, E. D. Abel, and M. E. Young. Disruption of the circadian clock within the cardiomyocyte influences myocardial contractile function, metabolism, and gene expression. American Journal of Physiology - Heart and Circulatory Physiology, 294(2):1036–1047, 2008.

[5] M. Brøns, M. Krupa, and M. Wechselberger. Mixed mode oscillations due to the generalized canard phenomenon. Fields Institute Communications, 49:39–63, 2006.

[6] C. R. Cederroth, U. Albrecht, J. Bass, S. A. Brown, J. Dyhrfjeld-Johnsen, F. Gachon, C. B. Green, M. H. Hastings, C. Helfrich-Förster, J. B. Hogenesch, F. Lévi, A. Loudon, G. B. Lundkvist, J. H. Meijer, M. Rosbash, J. S. Takahashi, M. Young, and B. Canlon. Medicine in the Fourth Dimension. Cell Metabolism, 30(2):238–250, 2019.

[7] M. G. Chang, D. Sato, E. De Lange, J. H. Lee, H. S. Karagueuzian, A. Garfinkel, J. N. Weiss, and Z. Qu. Bi-stable wave propagation and early afterdepolarization-mediated cardiac arrhythmias. Heart Rhythm, 9(1):115–122, 2012.

[8] S. L. Chellappa, N. Vujovic, J. S. Williams, and F. A. Scheer. Impact of Circadian Disruption on Cardiovascular Function and Disease. Trends in Endocrinology and Metabolism, 30(10):767–779, 2019.

[9] Y. Chen, D. Zhu, J. Yuan, Z. Han, Y. Wang, Z. Qian, and X. Hou. CLOCK-BMAL1 regulate the cardiac L-type calcium channel subunit CACNA1C through PI3K-Akt signaling pathway. Can. J. Physiol. Pharmacol., 94:1023–1032, 2016.

[10] D. J. Clemons and J. L. Seeman. The Laboratory Guinea Pig. CRC Press, 2nd edition, 2011.

[11] C. S. Colwell. Linking neural activity and molecular oscillations in the SCN. Nature Reviews Neuroscience, 12(10):553–69, 2011.

[12] E. De Lange, Y. Xie, and Z. Qu. Synchronization of early afterdepolarizations and arrhythmogenesis in heterogeneous cardiac tissue models. Biophysical Journal, 103(2):365–373, 2012.

[13] J. P. Degaute, E. Van Cauter, P. Van De Borne, and P. Linkowski. Twenty-four-hour blood pressure and heart rate profiles in humans: A twin study. Hypertension, 23(2):244–253, 1994.

[14] C. Diekman, M. Golubitsky, and T. McMillen. Reduction and Dynamics of a Generalized Rivalry Network. SIAM Journal on Applied Dynamical Systems, 11(4):1–33, 2012.

[15] P. E. Dilaveris, P. Färbom, V. Batchvarov, A. Ghuran, and M. Malik. Circadian behavior of P-wave duration, P-wave area, and PR interval in healthy subjects. Annals of Noninvasive Electrocardiology, 6(2):92–97, 2001.

[16] G. B. Ermentrout and D. H. Terman. Mathematical Foundations of Neuroscience. Springer, 2010.

[17] F. H. Fenton, E. M. Cherry, H. M. Hastings, and S. J. Evans. Multiple mechanisms of spiral wave breakup in a model of cardiac electrical activity. Chaos, 12(3):852–892, 2002.

[18] P. Ferdinandy, I. Baczkó, P. Bencsik, Z. Giricz, A. Görbe, P. Pacher, Z. V. Varga, A. Varró, and R. Schulz. Definition of hidden drug cardiotoxicity: Paradigm change in cardiac safety testing and its clinical implications. European Heart Journal, 40(22):1771–1777C, 2019.

[19] B. Fermini, J. C. Hancox, N. Abi-Gerges, M. Bridgland-Taylor, K. W. Chaudhary, T. Colatsky, K. Correll, W. Crumb, B. Damiano, G. Erdemli, G. Gintant, J. Imredy, J. Koerner, J. Kramer, P. Levesque, Z. Li, A. Lindqvist, C. A. Obejero-Paz, D. Rampe, K. Sawada, D. G. Strauss, and J. I. Vandenberg. A new perspective in the field of cardiac safety testing through the comprehensive in vitro proarrhythmia assay paradigm. Journal of Biomolecular Screening, 21(1):1–11, 2016.

[20] K. Fijorek, M. Puskulluoglu, and S. Polak. Circadian models of serum potassium, sodium, and calcium concentrations in healthy individuals and their application to cardiac electrophysiology simulations at individual level. Computational and Mathematical Methods in Medicine, 2013(429037):1–8, 2013.

[21] P. Fotiadis and D. B. Forger. Modeling the effects of the circadian clock on cardiac electrophysiology. Journal of Biological Rhythms, 28(1):69–78, 2013.

[22] A. Garfinkel, Y.-H. Kim, O. Voroshilovsky, Z. Qu, J. R. Kil, M.-H. Lee, H. S. Karagueuzian, J. N. Weiss, and P.-S. Chen. Preventing ventricular fibrillation by flattening cardiac restitution. Proceedings of the National Academy of Sciences, 97(11):6061–6066, 2000.

[23] L. A. Gottlieb, A. Lubberding, A. P. Larsen, and M. B. Thomsen. Circadian rhythm in QT interval is preserved in mice deficient of potassium channel interacting protein 2. Chronobiology International, 34(1):45–56, 2017.

[24] T. G. Granda, R. M. D’Attino, E. Filipski, P. Vrignaud, C. Garufi, E. Terzoli, M. C. Bissery, and F. Lévi. Circadian optimisation of irinotecan and oxaliplatin efficacy in mice with Glasgow osteosarcoma. British Journal of Cancer, 86(6):999–1005, 2002.

[25] S. Grubb, K. Calloe, and M. B. Thomsen. Impact of KChiP2 on cardiac electrophysiology and the progression of heart failure. Frontiers in Physiology, 3 MAY(May):1–9, 2012.

[26] P. Hammer. Spiral waves in monodomain reaction-diffusion model, 2008.

[27] R. C. Hillborn. Chaos and Nonlinear Dynamics: An Introduction for Scientists and Engineers. Oxford University Press, 2000.

[28] X. Huang, Z. Song, and Z. Qu. Determinants of early afterdepolarization properties in ventricular myocyte models. PLoS Computational Biology, 14(11):1–24, 2018.

[29] D. Jeyaraj, S. M. Haldar, X. Wan, M. D. McCauley, J. A. Ripperger, K. Hu, Y. Lu, B. L. Eapen, N. Sharma, E. Ficker, M. J. Cutler, J. Gulick, A. Sanbe, J. Robbins, S. Demolombe, R. V. Kondratov, S. A. Shea, U. Albrecht, X. H. Wehrens, D. S. Rosenbaum, and M. K. Jain. Circadian rhythms govern cardiac repolarization and arrhythmogenesis. Nature, 483(7387):96–101, 2012.

[30] B. S. Kim, Y. H. Kim, G. S. Hwang, H. N. Pak, S. C. Lee, W. J. Shim, D. J. Oh, and Y. M. Ro. Action potential duration restitution kinetics in human atrial fibrillation. Journal of the American College of Cardiology, 39(8):1329–1336, 2002.

[31] M. L. Ko, Y. Liu, S. E. Dryer, and G. Y. Ko. The expression of L-type voltage-gated calcium channels in retinal photoreceptors is under circadian control. Journal of Neurochemistry, 103(2):784–792, 2007.

[32] M. Kozák, L. Křivan, and B. Semrád. Circadian variations in the occurrence of ventricular tachyarrhythmias in patients with implantable cardioverter defibrillators. PACE - Pacing and Clinical Electrophysiology, 26(3):731–735, 2003.

[33] T. Krogh-Madsen and D. J. Christini. Nonlinear Dynamics in Cardiology. Annual Review of Biomedical Engineering, 14(1):179–203, 2012.

[34] P. Kügler. Early Afterdepolarizations with Growing Amplitudes via Delayed Subcritical Hopf Bifurcations and Unstable Manifolds of Saddle Foci in Cardiac Action Potential Dynamics. PLoS ONE, 11(3):30151178, 2016.

[35] P. Kügler, M. A. K. Bulelzai, and A. H. Erhardt. Period doubling cascades of limit cycles in cardiac action potential models as precursors to chaotic early Afterdepolarizations. BMC Systems Biology, pages 1–13, 2017.

[36] P. Kügler, A. H. Erhardt, and M. A. Bulelzai. Early afterdepolarizations in cardiac action potentials as mixed mode oscillations due to a folded node singularity. PLoS ONE, 13(12):1–22, 2018.

[37] Y. Kurata, K. Tsumoto, K. Hayashi, I. Hisatome, M. Tanida, Y. Kuda, and T. Shibamoto. Dynamical mechanisms of phase-2 early afterdepolarizations in human ventricular myocytes: insights from bifurcation analyses of two mathematical models. American Journal of Physiology - Heart and Circulatory Physiology, 312(1):H106–H127, 2017.

[38] K. N. Lee, S. T. Pellom, E. Oliver, and S. Chirwa. Characterization of the guinea pig animal model and subsequent comparison of the behavioral effects of selective dopaminergic drugs and methamphetamine. Synapse, 68(5):221–233, 2014.

[39] H. Li, W. Guo, R. L. Mellor, and J. M. Nerbonne. KChIP2 modulates the cell surface expression of Kv1.5-encoded K+ channels. Journal of Molecular and Cellular Cardiology, 39(1):121–132, 2005.

[40] Z. Li, G. R. Mirams, T. Yoshinaga, B. J. Ridder, X. Han, J. E. Chen, N. L. Stockbridge, T. A. Wisialowski, B. Damiano, S. Severi, P. Morissette, P. R. Kowey, M. Holbrook, G. Smith, R. L. Rasmusson, M. Liu, Z. Song, Z. Qu, D. J. Leishman, J. Steidl-Nichols, B. Rodriguez, A. Bueno-Orovio, X. Zhou, E. Passini, A. G. Edwards, S. Morotti, H. Ni, E. Grandi, C. E. Clancy, J. Vandenberg, A. Hill, M. Nakamura, T. Singer, L. Polonchuk, A. Greiter-Wilke, K. Wang, S. Nave, A. Fullerton, E. A. Sobie, M. Paci, F. Musuamba Tshinanu, and D. G. Strauss. General Principles for the Validation of Proarrhythmia Risk Prediction Models: An Extension of the CiPA In Silico Strategy [published online ahead of print November 10, 2019]. Clinical Pharmacology & Therapeutics, 2019.

[41] C. H. Luo and Y. Rudy. A model of the ventricular cardiac action potential. Depolarization, repolarization, and their interaction. Circulation Research, 68(6):1501–1526, 1991.

[42] R. V. Madhvani, M. Angelini, Y. Xie, A. Pantazis, S. Suriany, N. P. Borgstrom, A. Garfinkel, Z. Qu, J. N. Weiss, and R. Olcese. Targeting the late component of the cardiac L-type Ca2+ current to suppress early afterdepolarizations. Journal of General Physiology, 145(5):395–404, 2015.

[43] R. V. Madhvani, Y. Xie, A. Pantazis, A. Garfinkel, Z. Qu, J. N. Weiss, and R. Olcese. Shaping a new Ca 2+ conductance to suppress early afterdepolarizations in cardiac myocytes. Journal of Physiology, 589(24):6081–6092, 2011.

[44] Y. S. Markandeya and T. J. Kamp. Rational strategy to stop arrhythmias: Early afterdepolarizations and L-type Ca2+ current. Journal of General Physiology, 145(6):475–479, 2015.

[45] T. A. Martino and M. E. Young. Influence of the Cardiomyocyte Circadian Clock on Cardiac Physiology and Pathophysiology. Journal of Biological Rhythms, 30(3):183–205, 2015.

[46] B. McMillan, D. J. Gavaghan, and G. R. Mirams. Early afterdepolarisation tendency as a simulated pro-arrhythmic risk indicator. Toxicology Research, 6(6):912–921, 2017.

[47] J. E. Muller, P. L. Ludmer, S. N. Willich, G. H. Tofler, G. Aylmer, I. Klangos, and P. H. Stone. Circadian variation in the frequency of sudden cardiac death. Circulation, 75(1):131–138, 1987.

[48] S.-S. Nahm, Y. Z. Farnell, W. Griffith, and D. J. Earnest. Circadian Regulation and Function of Voltage-Dependent Calcium Channels in the Suprachiasmatic Nucleus. The Journal of Neuroscience, 25(40):9304–9308, 2005.

[49] M. Orini, N. Srinivasan, P. Taggart, and P. Lambiase. Reliability of APD-restitution slope measurements: Quantification and methodological comparison. Computing in Cardiology, 42:545–548, 2015.

[50] J. Parikh, P. Di Achille, J. Kozloski, and V. Gurev. Global sensitivity analysis of ventricular myocyte model-derived metrics for proarrhythmic risk assessment. Frontiers in Pharmacology, 10(October):1–18, 2019.

[51] C. M. Pennartz, M. T. De Jeu, N. P. Bos, J. Schaap, and A. M. Geurtsen. Diurnal modulation of pacemaker potentials and calcium current in the mammalian circadian clock. Nature, 416(6878):286–290, 2002.

[52] F. Portaluppi and R. C. Hermida. Circadian rhythms in cardiac arrhythmias and opportunities for their chronotherapy. Advanced Drug Delivery Reviews, 59(9-10):940–951, 2007.

[53] Z. Qu, L. H. Xie, R. Olcese, H. S. Karagueuzian, P. S. Chen, A. Garfinkel, and J. N. Weiss. Early afterdepolarizations in cardiac myocytes: Beyond reduced repolarization reserve. Cardiovascular Research, 99(1):6–15, 2013.

[54] B. M. D. Ruben, D. F. Smith, G. A. Fitzgerald, and J. B. Hogenesch. Dosing time matters. Science, 365(6453):547–550, 2019.

[55] M. D. Ruben, G. Wu, D. F. Smith, R. E. Schmidt, L. J. Francey, Y. Y. Lee, R. C. Anafi, and J. B. Hogenesch. A database of tissue-specific rhythmically expressed human genes has potential applications in circadian medicine. Sci Transl Med, 3(September):1–8, 2018.

[56] M. H. Ruwald, A. J. Moss, W. Zareba, C. Jons, A. C. Ruwald, S. McNitt, B. Polonsky, and V. Kutyifa. Circadian distribution of ventricular tachyarrhythmias and association with mortality in the MADIT-CRT trial. Journal of Cardiovascular Electrophysiology, 26(3):291–299, 2015.

[57] D. Sato, L.-h. Xie, T. P. Nguyen, J. N. Weiss, and Z. Qu. Irregularly Appearing Early Afterdepolarizations in Cardiac Myocytes: Random Fluctuations or Dynamical Chaos? Biophysical journal, 99(3):765–773, 2010.

[58] E. A. Schroder, D. E. Burgess, X. Zhang, M. Lefta, J. L. Smith, A. Patwardhan, D. C. Bartos, C. S. Elayi, K. A. Esser, and B. P. Delisle. The cardiomyocyte molecular clock regulates the circadian expression of Kcnh2 and contributes to ventricular repolarization. Heart Rhythm, 12(6):1306–1314, 2015.

[59] E. A. Schroder, M. Lefta, X. Zhang, D. C. Bartos, H.-z. Feng, Y. Zhao, A. Patwardhan, J.-p. Jin, K. A. Esser, and B. P. Delisle. The cardiomyocyte molecular clock, regulation of Scn5a, and arrhythmia susceptibility. Am J Physiol Cell Physiol, pages 954–965, 2013.

[60] P. Seenivasan, S. N. Menon, S. Sridhar, and S. Sinha. When the clock strikes : Modeling the relation between circadian rhythms and cardiac arrhythmias. Journal of Physics: Conference Series, 759(012021):1–8, 2016.

[61] P. S. Spector. Diagnosis and management of sudden cardiac death. Heart, 91(3):408–413, 2005.

[62] E. C. Stecker, K. Reinier, E. Marijon, K. Narayanan, C. Teodorescu, A. Uy-Evanado, K. Gunson, J. Jui, and S. S. Chugh. Public health burden of sudden cardiac death in the United States. Circulation: Arrhythmia and Electrophysiology, 7(2):212–217, 2014.

[63] R. K. Thakur, R. G. Hoffmann, D. W. Olson, R. Joshi, D. D. Tresch, T. P. Aufderheide, and J. H. Ip. Circadian variation in sudden cardiac death: Effects of age, sex, and initial cardiac rhythm. Annals of Emergency Medicine, 27(1):29–34, 1996.

[64] M. Tong, S. Wang, Y. Pang, Y. Zhou, H. Cui, L. Ruan, J. Su, and X. Chen. Circadian expression of connexins in the mouse heart. Biological Rhythm Research, 47(4):631–639, 2016.

[65] M. Tong, E. Watanabe, N. Yamamoto, M. Nagahata-Ishiguro, K. Maemura, N. Takeda, R. Nagai, and Y. Ozaki. Circadian expressions of cardiac ion channel genes in mouse might be associated with the central clock in the SCN but not the peripheral clock in the heart. Biological Rhythm Research, 44(4):519–530, 2013.

[66] D. X. Tran, D. Sato, A. Yochelis, J. N. Weiss, A. Garfinkel, and Z. Qu. Bifurcation and chaos in a model of cardiac early afterdepolarizations. Physical Review Letters, 102(25):1–4, 2009.

[67] N. Vandersickel, T. P. D. Boer, M. A. Vos, and A. V. Panfilov. Perpetuation of torsade de pointes in heterogeneous hearts : competing foci or re-entry? J Physiol, 23:6865–6878, 2016.

[68] N. Vandersickel, I. V. Kazbanov, A. Nuitermans, L. D. Weise, R. Pandit, and A. V. Panfilov. A study of early afterdepolarizations in a model for human ventricular tissue. PLoS ONE, 9(1):e84595, 2014.

[69] P. C. Viswanathan, R. M. Shaw, and Y. Rudy. Effects of IKr and IKs Heterogeneity on Action Potential Duration and Its Rate Dependence: A Simulation Study. Circulation, 99:2466–2474, 1999.

[70] T. Vo and R. Bertram. Why pacing frequency affects the production of early afterdepolarizations in cardiomyocytes: An explanation revealed by slow-fast analysis of a minimal model. Physical Review E, 99(5):52205, 2019.

[71] J. N. Weiss, A. Garfinkel, H. S. Karagueuzian, P. S. Chen, and Z. Qu. Early afterdepolarizations and cardiac arrhythmias. Heart Rhythm, 7(12):1891–1899, 2010.

[72] S. N. Willich, D. Levy, M. B. Rocco, G. H. Tofler, P. H. Stone, and J. E. Muller. Circadian variation in the incidence of sudden cardiac death in the framingham heart study population. The American Journal of Cardiology, 60(10):801–806, 1987.

[73] Y. Xie, D. Sato, A. Garfinkel, Z. Qu, and J. N. Weiss. So little source, so much sink: Requirements for afterdepolarizations to propagate in tissue. Biophysical Journal, 99(5):1408–1415, 2010.

[74] T. Yamashita, A. Sekiguchi, Y. ki Iwasaki, K. Sagara, H. Iinuma, S. Hatano, L. T. Fu, and H. Watanabe. Circadian variation of cardiac K+ channel gene expression. Circulation, 107(14):1917–1922, 2003.

[75] M. E. Young, R. A. Brewer, R. A. Peliciari-Garcia, H. E. Collins, L. He, T. L. Birky, B. W. Peden, E. G. Thompson, B. J. Ammons, M. S. Bray, J. C. Chatham, A. R. Wende, Q. Yang, C. W. Chow, T. A. Martino, and K. L. Gamble. Cardiomyocyte-specific BMAL1 plays critical roles in metabolism, signaling, and maintenance of contractile function of the heart. Journal of Biological Rhythms, 29(4):257–276, 2014.

[76] J. Zhang, J. Chatham, and M. E. Young. Circadian Regulation of Cardiac Physiology: Rhythms That Keep the Heart Beating. Annual Review of Physiology, 82(1):1–23, 2020.

[77] Z. Zhao, Y. Xie, H. Wen, D. Xiao, C. Allen, N. Fefelova, W. Dun, P. A. Boyden, Z. Qu, and L.-H. Xie. Role of the transient outward potassium current in the genesis of early afterdepolarizations in cardiac cells. Cardiovascular Research, 95:308–316, 2012.

